# Extending Kilosort for 3D Neural Probes

**DOI:** 10.64898/2026.05.14.725070

**Authors:** Jia-Hao Zhang, Jyh-Jang Sun, Kuan-Peng Chen, Kuo-Hsing Kao, Nan-Yow Chen

## Abstract

Kilosort 2.0 is a widely adopted spike sorting algorithm recognized for its efficiency and accuracy on planar electrode arrays, such as Neuropixels. To adapt its robust architecture to emerging three-dimensional (3D) neural probes, we present Kilosort 2.0-3D, a modified version that leverages 3D spatial information. Our modification specifically redefines the spatial processing components of Kilosort 2.0 to operate in 3D space while leaving the core template-matching process unchanged. By using synthetic extracellular recording data with ground-truth neuron positions and firing times, we demonstrate that Kilosort 2.0-3D effectively resolves spatial ambiguities and unit misclassifications inherent in 2D spatial assumptions. Our results show that Kilosort 2.0-3D achieves rotational invariance and maintains full backward compatibility with planar arrays. This work establishes a validated, scalable tool for spike sorting of high-density 3D neural electrophysiology data.

## Introduction

Simultaneously monitoring thousands of neurons has become a cornerstone of modern extracellular electrophysiology (Buzsáki, 2004; Steinmetz et al., 2019). A crucial step in this pipeline is spike sorting, the process of identifying individual neurons from multichannel extracellular signals. Driven by the need for higher-precision monitoring and more accurate spatial localization, neural probe technology has evolved from microwires and tetrodes (Gray et al., 1995) to high-density silicon arrays (Jun et al., 2017; Steinmetz et al., 2021), and more recently, toward three-dimensional (3D) architectures (Obaid et al., 2020; Sahasrabuddhe et al., 2021; Musk et al., 2019; Yang et al., 2025). These emerging 3D configurations offer significant advantages such as improved spatial resolution and better motion compensation (Boussard et al., 2021). Furthermore, their volumetric design allows researchers to move beyond the constraints of planar probes, enabling simultaneous sampling of large and diverse populations of neurons across both vertical cortical layers and lateral functional domains. This capability is crucial for investigating how spatially distributed neural circuits perform complex computations (Senzai et al., 2019).

As recording hardware advances, spike sorting algorithms must be updated accordingly. Among existing tools, Kilosort 2.0 is one of the most widely adopted frameworks, well known for its speed, accuracy, and strong performance on planar probes (Pachitariu et al., 2016; Steinmetz et al., 2021). Its success largely derives from a powerful template-matching algorithm that is based on the 2D high-density electrode geometries. However, this reliance poses a limitation. When applied directly to true 3D configurations, the algorithm inherently forces a geometric projection onto a 2D plane. This dimensional collapse misrepresents the true Euclidean inter-channel distances, leading to severe spatial aliasing, artificial merging of distinct units, and ultimately degraded sorting performance.

To address this, we introduce Kilosort 2.0-3D, a targeted extension of the Kilosort 2.0 (with GNU General Public License) framework for 3D configurations. Rather than developing a new framework from scratch, our approach extends the existing and widely trusted Kilosort 2.0 framework by adapting its spatial metric to operate directly in 3D Euclidean space. This preserves its efficient core while ensuring seamless integration into established analytical workflows. Using synthetic extracellular datasets that mimic ideal volumetric recordings, we demonstrate that a true 3D spatial model is essential for accurate sorting. We show that our extension effectively recovers neural units, achieves rotational invariance, and maintains full backward compatibility with planar probes. Together, these results establish a geometry-agnostic extension of Kilosort 2.0, providing a reliable tool for next-generation neural recording systems.

## Methods

### Modification of Kilosort for 3D Electrode Arrays

The Kilosort 2.0 pipeline relies on the spatial relationships between recording channels to define local neighborhoods for spatiotemporal template matching. To adapt the algorithm for 3D probe geometries, we implemented two key modifications, as conceptually illustrated in Fig. 1.

**Fig. 1.**
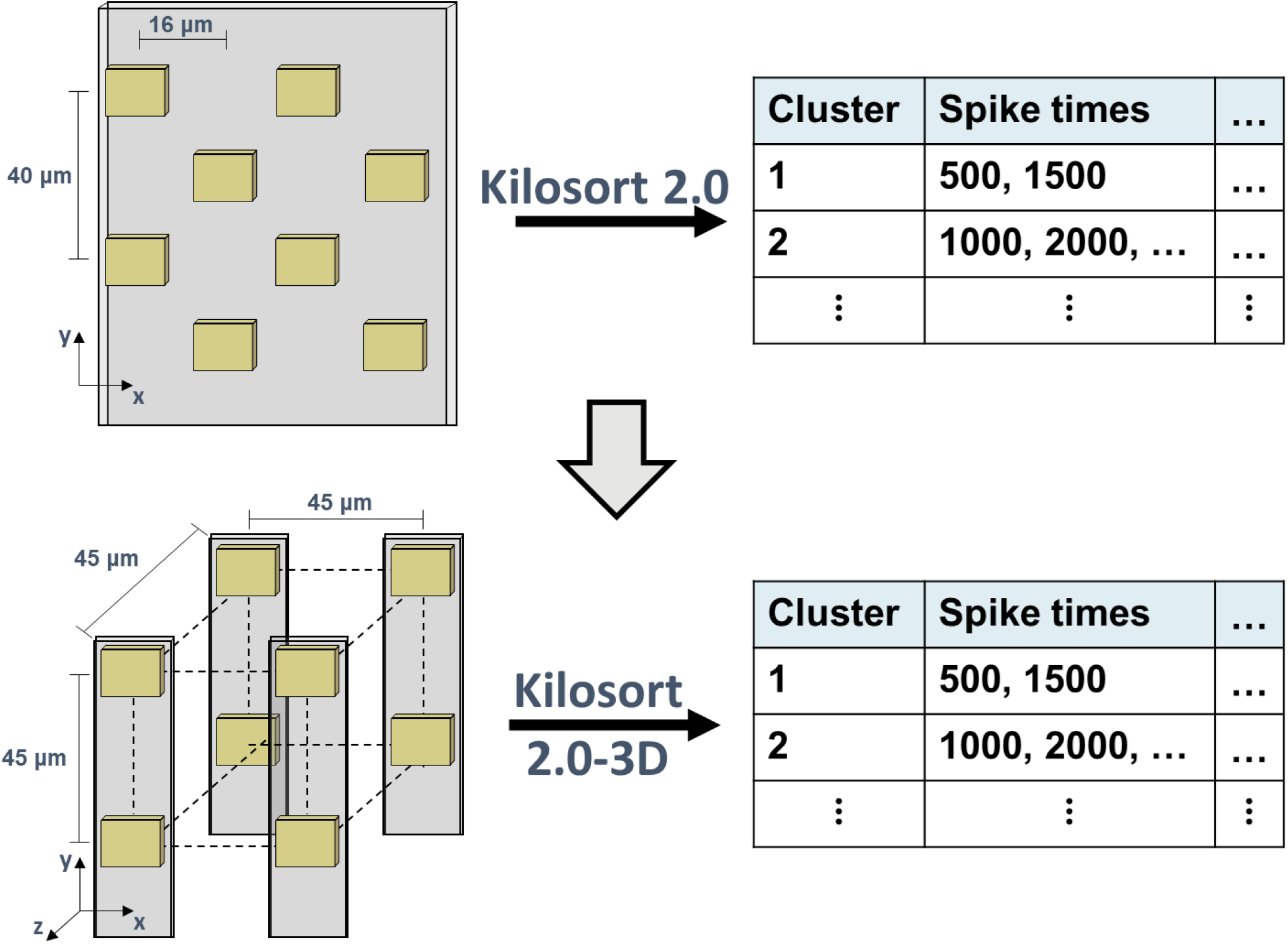
Schematic overview of the Kilosort 2.0 extension for 3D neural probes. (Top) Standard Kilosort 2.0 framework utilizes a 2D channel map. (Bottom) The proposed Kilosort 2.0-3D framework integrates the whole 3D channel map, enabling utilization of full 3D coordinates during distance calculation. Both frameworks retain the same output structure, ensuring compatibility with existing downstream analysis workflows.

First, the channel map loading routine was updated to accept three-dimensional coordinates for each recording site. For planar probes, this is accommodated by setting the new z-coordinate to a constant value (e.g., zero). Second, and most critically, the internal distance calculation function, i.e. getClosestChannels.m, was modified to compute the full 3D Euclidean distance between channels, rather than the original 2D projection:

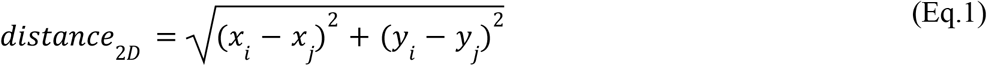

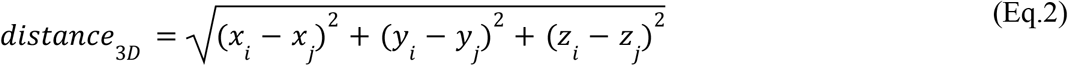

Where (*x*_*i*_, *y*_*i*_, *z*_*i*_) and (*x*_*j*_, *y*_*j*_, *z*_*j*_) denote the Euclidean coordinates of channels *i* and *j*, respectively. The core components of the algorithm, including template matching, clustering, and drift correction, were intentionally left unchanged to preserve the proven performance characteristics of the original Kilosort 2.0.

### Synthetic Dataset Generation

To rigorously validate our Kilosort 2.0-3D extension, we performed a series of in silico extracellular simulations that provided a ground-truth reference for spike sorting. A total of 20 independent datasets were synthesized, each encompassing a defined 3D volume of 894×894×3970 μm^3^, symmetrically distributed around the origin along all three axes (447 μm in x and z, 1985 μm in y for one side) (Yang et al., 2025). This space contained 1000 rat cortical neurons and one of three probe models: a standard planar Neuropixels probe, or a 384-channel Simple Cubic probe arranged in either a 4×4×24 cubic or a 2×2×96 column. Both SC probes featured a uniform 45 μm inter-electrode spacing. Notably, the 2×2×96 geometry was specifically designed as a reference with the Neuropixels probe. Each probe was centered at the origin, with neuronal placement constrained to prevent physical overlap with the probe structure.

As depicted in Fig. 2, neurons were distributed across six cortical layers, beginning at y =1000 μm and extending downward along the y-axis. The thicknesses of these layers were set to 157, 418, 325, 511, and 562 μm respectively (Narayanan et al., 2017). Notably, layers 2 and 3 were consolidated into a single layer due to similarities in neuronal composition and an indistinct boundary. Neurons were assigned to the layers in proportions of 0.0019, 0.2177, 0.2753, 0.2168, and 0.2883 (Narayanan et al., 2017), respectively. Within each layer, neuron positions were sampled uniformly at random. To avoid overlap, no two neurons were placed within 10 μm of each other, and all neurons were positioned at least 10 μm away from the probe.

**Fig. 2.**
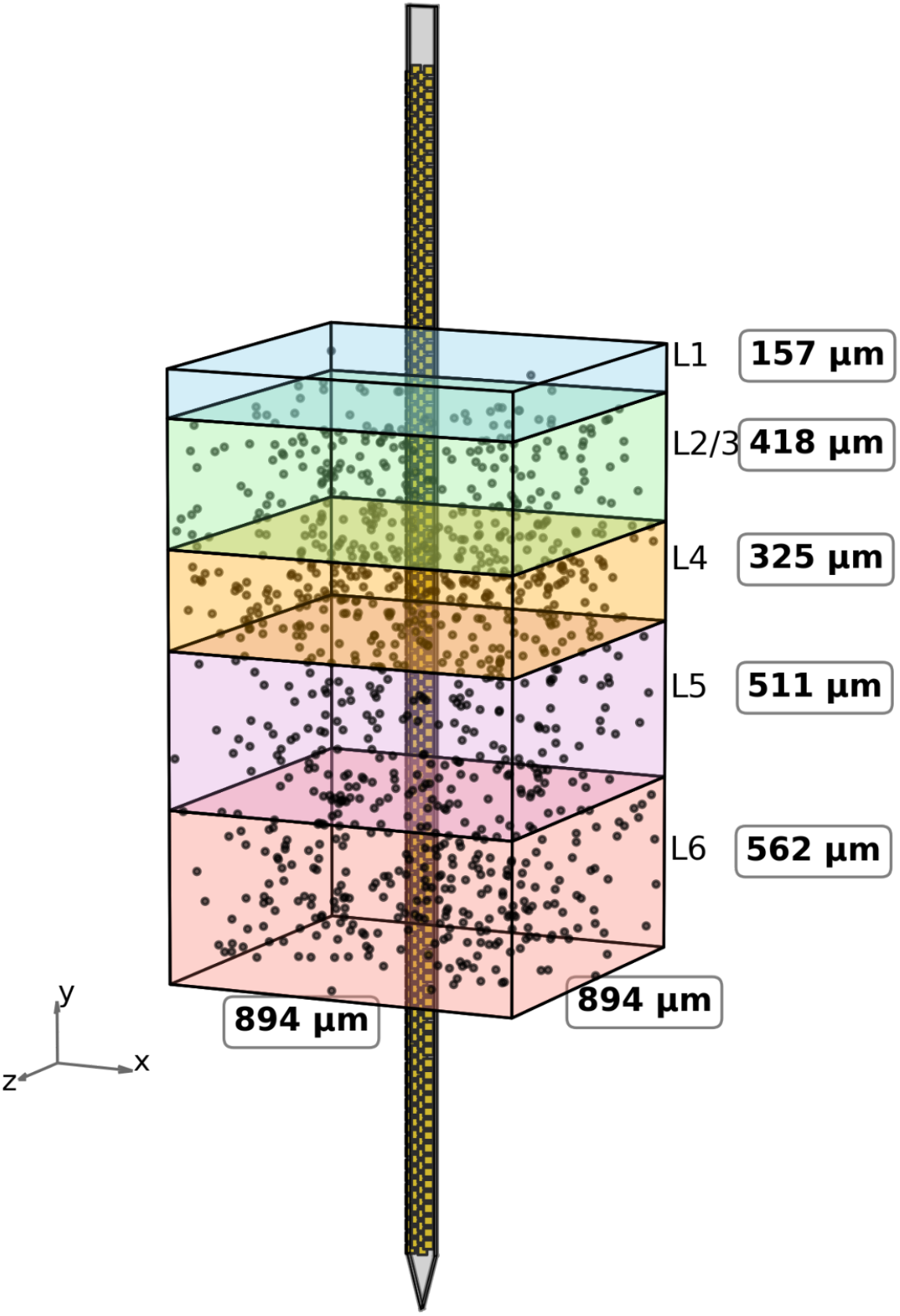
Distribution of cortical neurons surrounding a Neuropixels probe. A color-coded cortical column is divided into five layers (L1, L2/3, L4, L5, L6) with neurons (black dots) positioned according to layer-specific densities. The probe (gray) penetrates vertically through all layers, with its center aligned at the origin.

In silico extracellular potentials were synthesized using the NEURON (Hines et al., 2009) and LFPy (Hagen et al., 2018) simulation toolkits, enabling accurate computation of extracellular signals and their contributions to local field potentials at the recording sites. Neuronal morphology and electrophysiological properties were sourced from the Neocortical Microcircuit Collaboration Portal of the Blue Brain Project (Markram et al., 2015). The dataset includes 207 morpho-electrical types, combined by 55 different morphological types with 11 electrophysiological types. Identical neuronal configurations were maintained across all probe geometries to ensure fair and controlled comparisons of spike sorting performance between the Kilosort 2.0-3D and the original algorithm.

### Evaluation Setup

We designed three distinct tests to comprehensively evaluate the performance, robustness, and backward compatibility of Kilosort 2.0 and Kilosort 2.0-3D. The output of each sorting session was evaluated using custom tools designed to assess cluster separation and quality.

#### Environment Validation

To validate the reliability of our synthetic datasets, we utilized a standard planar Neuropixels probe within the original Kilosort 2.0 framework. This procedure ensures that the simulation environment, including neuronal densities and extracellular potential models, is appropriately configured to provide a robust and credible baseline for subsequent algorithmic testing.

#### Rotational Invariance

To verify geometric robustness, we applied various rigid rotations to the entire 3D dataset (affecting both probe and neuron coordinates) before processing with Kilosort 2.0-3D. We required the sorting output to be consistent with the results from the unrotated dataset, confirming the stability of the 3D Euclidean metric.

#### Backward Compatibility

To ensure that our modifications did not impair performance on conventional probe geometries, we processed synthetic datasets generated for a standard Neuropixels probe. Both the original Kilosort 2.0 and Kilosort 2.0-3D (with z-coordinates set to zero) were run on this 2D data to confirm that they produced identical sorting results.

### Performance Metrics

To rigorously quantify the sorting outputs across all 20 synthesis datasets, we evaluated the results based on three standard metrics: True Positives (TP), False Positives (FP), and False Negatives (FN).

#### True Positive (TP)

Defined as an identified cluster that uniquely (one-to-one) matches the identity of a single ground-truth neuron, with spike timings aligned within a 1 ms temporal tolerance.

#### False Positive (FP)

Defined as an identified cluster that cannot entirely correspond to any ground-truth neuron, often resulting from noise artifacts or the erroneous over-splitting of a single actual neuron.

#### False Negative (FN)

Occurs when a ground-truth neuron fails to be uniquely identified by the algorithm, typically due to missed detection or merging with similar templates.

### Discovery Volume and Normalization

To quantify the spatial coverage and facilitate performance comparisons across diverse 3D probe configurations, the following metrics were implemented

#### Discovery Volume

To quantify the spatial reach of each probe design, we defined the discovery volume as the total union of spherical spaces centered at each electrode’s coordinates with a discovery radius of 75 μm. This radius represents the empirical detection range for the recording system, as validated by Yang et al. (2025). By accounting for spatial overlaps between neighboring electrodes, this metric provides a more accurate representation of the three-dimensional tissue coverage than simple lateral area.

#### Effective Electrodes and Metric Normalization

Effective electrodes are defined as those having at least one ground-truth neuron within their 75 μm radius. This definition ensures that electrodes positioned in empty neural space do not artificially reduce the performance. To ensure a fair and rigorous comparison between probe designs with varying electrode counts, all reported performance metrics were normalized by the number of effective electrodes.

### Implementation of Kilosort 2.0-3D

To support the analysis of 3D neural recordings, we systematically extended the original Kilosort 2.0 framework (MATLAB-based) to incorporate the z-dimension. The Kilosort 2.0-3D focused on upgrading spatial distance metrics, data structures, and preprocessing routines to accommodate 3D coordinates. A summary of the key code modifications is provided in Table 1.

**Table 1.** Summary of Kilosort 2.0-3D Code Modifications.

| File | Kilosort 2.0 (original ) | Kilosort 2.0-3D (This Work) | Functional Impact |
| --- | --- | --- | --- |
| getClosestChannels.m<br>whiteningLocal.m | $(\text{rez.xc}(:) - \text{rez.xc}(:))^2 + (\text{rez.yc}(:) - \text{rez.yc}(:))^2$ | $(\text{rez.xc}(:) - \text{rez.xc}(:))^2 + (\text{rez.yc}(:) - \text{rez.yc}(:))^2 + (\text{rez.zc}(:) - \text{rez.zc}(:))^2$ | Enables full 3D distance calculation. |
| preprocessDataSub.m<br>loadChanMap.m<br>get_whitening_matrix.m | $\text{xc} = \text{rez.xc}$<br>$\text{yc} = \text{rez.yc}$ | $\text{xc} = \text{rez.xc}$<br>$\text{yc} = \text{rez.yc}$<br>$\text{zc} = \text{rez.zc}$ | Ensures z-axis information persists through. |

## Results and Analysis

### Maintained Backward Compatibility with Planar Probes

A fundamental requirement for our algorithmic extension is that it must not compromise performance on conventional recording tasks. To verify this, we evaluated backward compatibility using a synthetic dataset and recorded using a planar Neuropixels probe. A 2D channel map was processed using both the original Kilosort 2.0 and Kilosort 2.0-3D. As illustrated in the first and second columns of Fig. 3 (**NPX + Kilosort 2.0 and NPX + Kilosort 2.0-3D**), the sorting outputs were numerically identical. We observed a perfect one-to-one correspondence for all identified units and their respective spike times. This result confirms that our modifications to the spatial metric do not affect 2D channel map operations, ensuring that Kilosort 2.0-3D can be safely deployed on standard planar array data without any loss of fidelity.

**Fig. 3.**
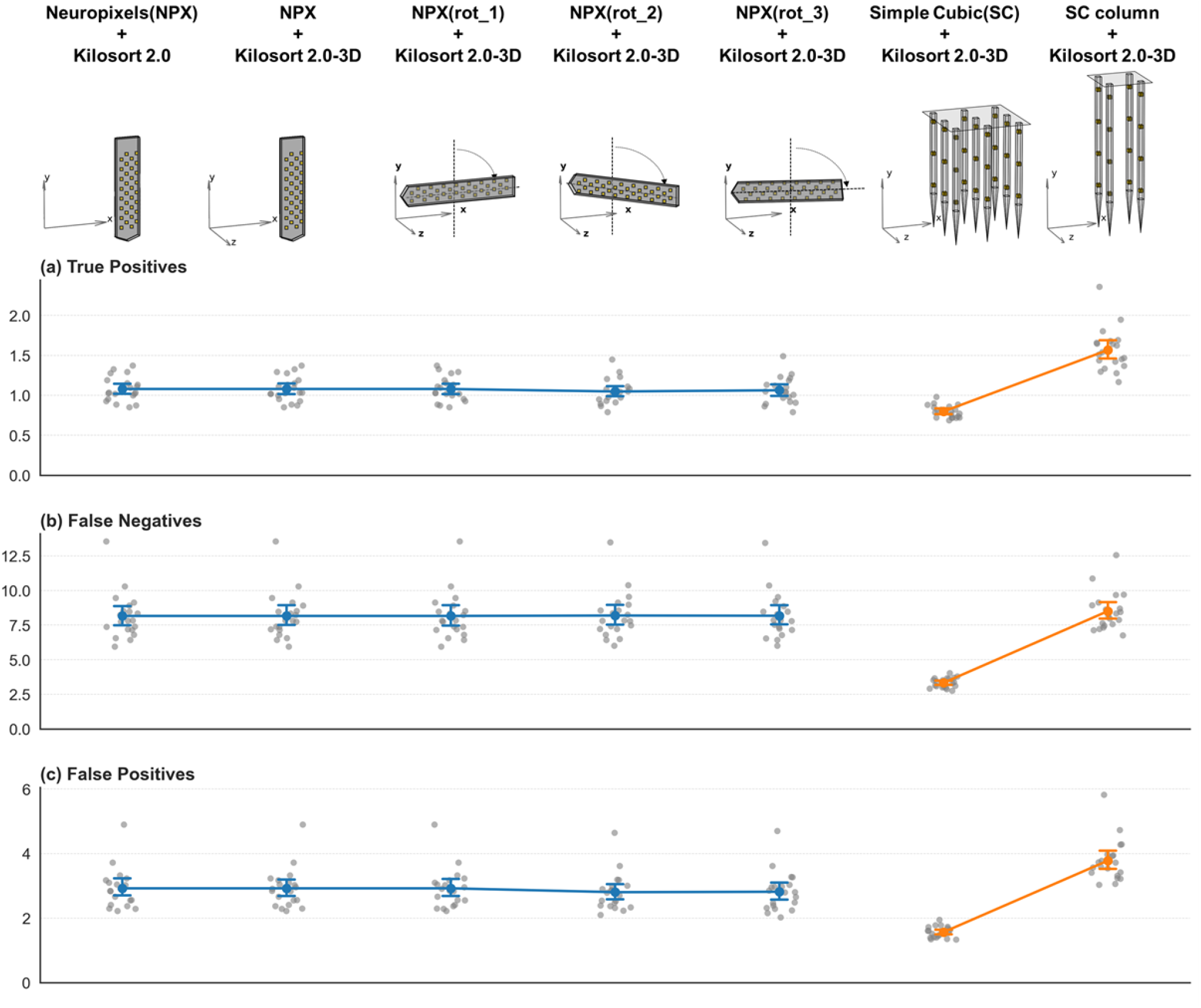
Performance evaluation of the modified Kilosort 2.0-3D algorithm across various probe geometries and spatial orientations (SNR=0dB). **(a-c)** Spike sorting metrics: (a) True Positives, (b) False Negatives, and (c) False Positives. All y-axis values are normalized by the number of effective electrodes (defined as electrodes having at least one neuron within 75 μm radius). Results using the planar Neuropixels (NPX) probe with different status to confirm the dataset working and that Kilosort 2.0-3D maintains full backward compatibility and rotational invariance. In 3D configurations, the Simple Cubic (SC) column, which has a similar distribution to NPX, leverages its 3D layout for improved yield. Conversely, the SC cubic exhibits the lowest normalized values due to high spatial redundancy; its grid-like structure increases the count of effective electrodes per neuron, thereby raising the normalization denominator and resulting in a more conservative efficiency profile.

### Rotational Invariance of the Kilosort 2.0-3D

To ensure the algorithm is robust to variations in physical probe orientation, we conducted a rigorous test for rotational invariance. The entire synthetic dataset, encompassing both probe and neuron coordinates, was subjected to various rigid 3D rotations prior to processing. As shown in the third to fifth columns of Fig. 3 (**NPX with rotation 1, 2, 3 + Kilosort 2.0-3D**), Kilosort 2.0-3D produced sorting outputs that were highly consistent with the results from the original, un-rotated configuration.

The minor discrepancies observed are primarily attributed to floating-point precision approximations and discretization artifacts introduced during the mathematical rotation of coordinates. Because high-density probes possess a discrete and symmetric geometric structure, these infinitesimal numerical shifts can occasionally influence the selection of adjacent channel neighborhoods during the template-matching phase, leading to subtle variations in final clustering. Despite these precision-induced fluctuations, the overall sorting performance remained stable, demonstrating that our 3D Euclidean distance calculations are geometrically sound and fundamentally independent of the probe’s orientation in space.

### Resolution of Spatial Ambiguity

The core advantage of Kilosort 2.0-3D is resolving spatial ambiguities that are fundamentally intractable in 2D. As illustrated in Fig. 4 (Left), when two neurons are mirrored across a planar probe surface, their signals superimpose on identical electrode sets. Lacking spatial diversity, the original Kilosort 2.0 fails to separate them, leading to a “false Merge” into a single contaminated cluster.

**Fig. 4.**
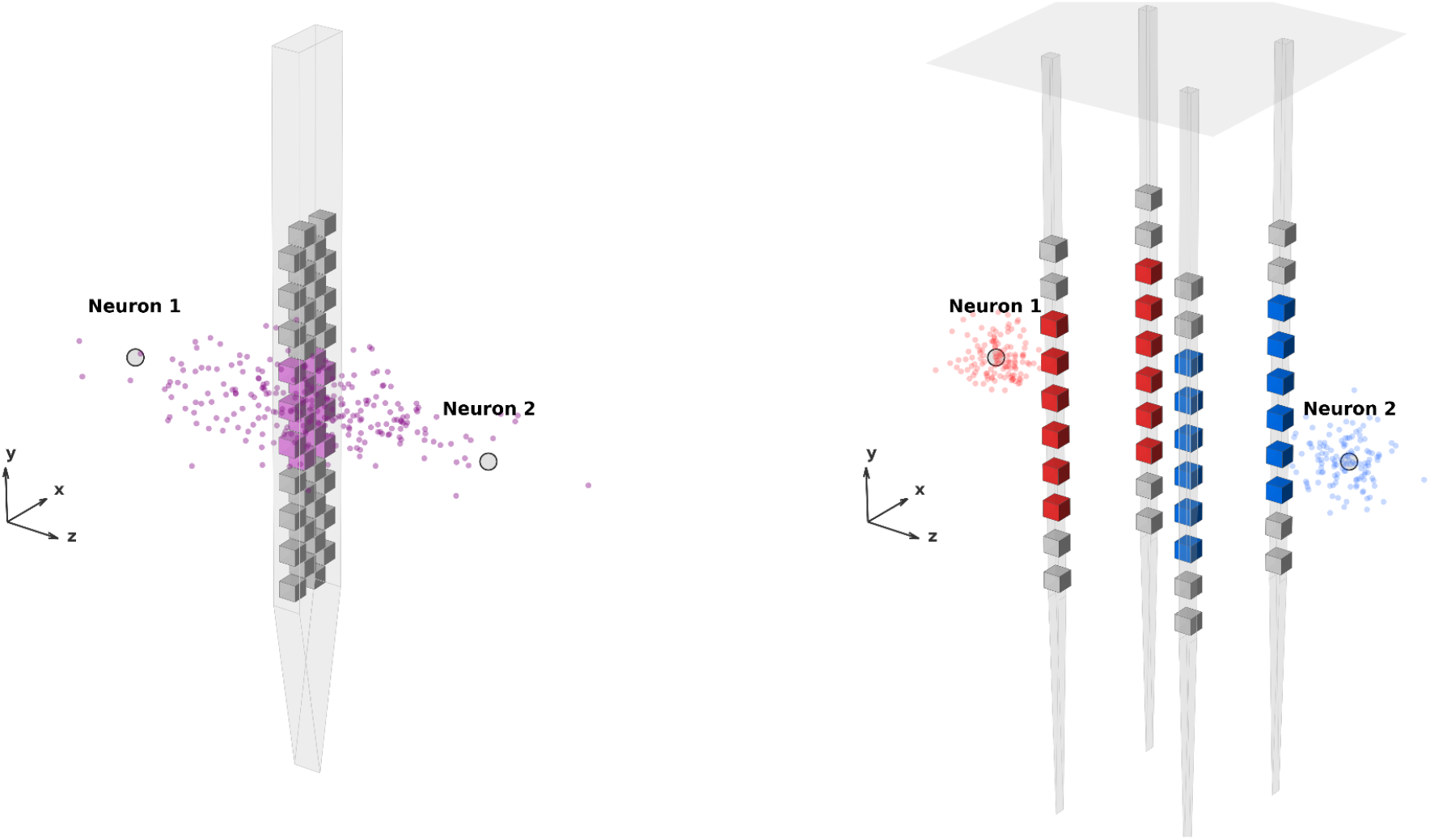
Resolution of Spatial Ambiguity. **(Left) Planar Probe Constraints:** Using a standard Neuropixels probe, the system lacks detection orthogonal to the probe surface. Regardless of the sorting algorithm used, the signal footprints of two distinct neurons superimpose, causing a “false merge” into a single contaminated cluster (purple). **(Right) 3D Advantage (3D Probe + Kilosort 2.0-3D):** By utilizing a 3D probe geometry coupled with the Kilosort 2.0-3D, the system leverages full 3D geometry. This combination successfully resolves the spatial offset, correctly separating the previously merged cluster into two distinct ground-truth units (red and blue).

However, by pairing Kilosort 2.0-3D with 3D probes (Fig. 4, Right), the system captures signals from multiple orientations to improve spatial resolution. This allows the algorithm to utilize distinct electrode combinations to distinguish units even when they appear identical on a planar surface. This hardware-software synergy is clearly reflected in the superior performance of the Simple Cubic (SC) column shown in Fig. 3. By leveraging its 3D layout to resolve these ambiguities, the Simple Cubic (SC) column achieves significantly higher normalized true positive counts (1.57 ± 0.06) compared to planar models (1.07 ± 0.02; t-test, *p* < 0.001).

### The Impact of Spatial Redundancy

Although it has been suggested that 3D probe designs should prioritize expanding the lateral direction to maximize the discovery volume and capture a larger population of neurons (Yang et al., 2025), our data reveals a contradictory trend. Contrary to the expectation that a larger discovery area captures more neurons, the SC cubic (4×4×24), which has the largest lateral coverage, consistently exhibits the lowest performance metrics (1.18 ± 0.07) compared to SC column (2×2×96) (1.57 ± 0.06; t-test, *p* < 0.01) as shown in Fig. 3.

This discrepancy indicates that the benefits of a larger discovery volume are scale-dependent. At a 45 μm inter-electrode spacing, the expanded grid of the cubic introduces an extreme redundancy penalty. Instead of providing unique spatial information, the dense arrangement of electrodes causes a single neuron’s signal to be captured by an excessive number of redundant channels. This results in a massive inflation of the effective electrodes count (the normalization denominator), which far outweighs any potential gains in unit detection.

Therefore, our results demonstrate that at high densities, the “wider view” of a cubic geometry actually becomes a shortage. While such designs may excel at larger scales (e.g., 150 μm), they suffer from severe efficiency degradation at 45 μm due to signal overlap and redundancy. In this high-density regime, the SC column proves to be the superior configuration. It provides sufficient 3D spatial diversity to resolve ambiguities while maintaining a leaner electrode profile that avoids the redundancy trap inherent in the cubic design.

## Discussion and Conclusion

In this study, we extended the Kilosort 2.0 algorithm for 3D electrode arrays through a minimal yet critical modification of its core distance metric. By analyzing 20 synthetic datasets, we successfully addressed the “Dimensional Collapse” inherent in planar assumptions, demonstrating that this 3D-aware spatial processing is essential for accurately resolving neural signals within volumetric recordings.

Furthermore, this study reveals that the trade-off between discovery volume and spatial redundancy is more complex than previously understood. While previous work (Yang et. al, 2025) emphasized the advantages of wide-area sampling at larger electrode spacings, our results demonstrate that these benefits are not unconditional. Specifically, expanding the recording volume only enhances performance when electrode density is maintained at a sufficiently low level. In high-density configurations, further lateral expansion, despite yielding the highest discovery volume, introduces extreme spatial redundancy that outweighs any gain in signal coverage. This suggests that “seeing more” is only beneficial if the information captured remains unique. Consequently, our findings complement existing literature by supporting a “sparse-optimized” probe layout principle. This suggests that discovery volume should be optimized rather than maximized. At high densities, redundant electrodes merely inflate the normalization denominator without providing unique neural information, thereby diluting overall sorting efficiency.

The primary value of this tool lies in its role as a critical bridge between hardware development and software analysis. As 3D probe technologies evolve, the research community requires a stable and trusted analysis platform that does not force changes to established analytical workflows. Kilosort 2.0-3D preserves the efficient architecture of the original while offering a geometry-agnostic design that allows researchers to directly evaluate the performance of various 3D layouts. Regarding stability, the algorithm demonstrates robust rotational invariance and full backward compatibility. Whether processing legacy planar data or 3D experiments with complex insertion angles, ensuring that results remain highly consistent whether processing legacy planar data or modern 3D experiments with complex surgical insertion angles.

Moreover, the Kilosort 2.0-3D logic developed in this work is architecturally aligned with the newly released Kilosort 4. While Kilosort 4 transitions to a Python-native, graph-based clustering algorithm, it retains the fundamental spatiotemporal template matching framework that shares the same core of our 3D extension. As detailed in our Methods section, the 3D coordinate mapping and distance calculation are designed to be directly transferable, ensuring that the community can transition to a “Kilosort 4-3D” pipeline with minimal integration overhead.

In conclusion, Kilosort 2.0-3D provides a robust and forward-looking extension that bridges the gap between hardware innovation and signal processing, establishing the necessary technical infrastructure for the era of high-resolution volumetric brain circuit studies.

## Code Availability

The Kilosort 2.0-3D code used in this study is available from the corresponding author upon reasonable request. We are in the process of preparing a public release (GNU General Public License) on a GitHub repository.

## Acknowledgements

We thank the National Center for High-performance Computing (NCHC) for providing computational and storage resources that facilitated this research.

